# Generation and quality control of maternal plasma lipidomics data associated with preterm birth

**DOI:** 10.1101/714790

**Authors:** ZhanLong Mei, Lingfei Ye, Kang Huang, Xi Yang, Xiaomin Chen, Miaolan Cen, Yuan Chen, Sujun Zhu, Juan Zeng, Bhaskar Roy, Hui Jiang, Wen-Jing Wang

## Abstract

Preterm birth is not only one of the most common causes of infant deaths but also a great risk for them to have severe subsequent health problems. The causes of preterm birth may be due to a combination of genetic and environmental factors, however, it remains largely unknown. Here we report an untargeted lipidomics dataset of plasma specimens from 258 pregnant women at the stage of twelve to twenty-five gestational weeks. Among them, 44 had extremely to very preterm births, 54 had moderate preterm births, 71 had late preterm births and 89 had full-term deliveries. The metabolomic profiling was generated with an UPLC-MS in both the positive and negative mode, and putative identification of all the metabolites was provided by searching against online databases. The quality assessment performed on quality control samples showed that the data is reproducible, robust and reliable. Both the raw data files, the raw and processed data matrix were available on MetaboLights, which may be used as a valuable validation dataset for new findings and a test dataset for novel algorithms.

## Background & Summary

Preterm birth (PTB) refers to delivery before 37 completed weeks of gestation, and it is one of the “Great Obstetrical Syndromes” ^1^. The global incidence of PTB was 9.6%, and more than 1.17 million PTB happen in China each year, making the second largest PTB population ^2^. PTB is the leading cause of neonatal mortality and morbidity, and more than one million babies die each year from PTB complications ^2^. Premature babies have an increased chance to have complications such as breathing difficulties, underdeveloped organs, low birth weight, cerebral palsy, and vision or hearing problems. According to the gestational age of delivery, PTB is classified as extremely preterm (<28 weeks), very preterm (28-<32 weeks) and moderate preterm (32-<34 weeks) or late preterm (34-<37 weeks). The earlier the delivery happens, the more adverse the consequence is.

The cause of PTB is complicated, and it could result from infection, inflammation, stress, genetic factors or induced for medical reasons ^3,4^. Once preterm labor started, there is no sure way to stop it, but medical treatments could help to slow down the procedure and reduce the risk of premature birth complications. Although there are many tests using cervicovaginal fetal fibronectin (fFN) or transvaginal sonographic cervical length measurements to predict the risk of PTB, accurate prediction of PTB remains a challenging task ^5^.

Lipids serving as fuel molecules, signal molecules and components of membranes components are important to pregnancy. There is a close relationship between PTB and progesterone, and weekly injection of 17 alpha-hydroxyprogesterone caproate could prevent recurrent preterm delivery ^6,7^. As the precursor of steroid hormones, maternal cholesterol concentration and its association with PTB has been investigated in many studies, and the results are controversial ^8^. The role of omega-3 (n-3) long-chain polyunsaturated fatty acids (LCPUFA) in pregnancy has also been studied, and cervical ripening and uterine contractions could be affected by the local concentration of specific fatty acids ^9–11^. Randomized trials of total LCPUFA supplementation have been conducted and found a reduction in early PTB (<34 weeks) ^10,12,13^. The second-trimester serum lipid profiling of 35 PTB women and 35 matched term delivery controls were carried out, and increased levels of lipids and changes of key end-point metabolites were found in PTB women ^14^. All these discoveries imply that lipids could play an important role in PTB and could be used as potential biomarkers for PTB or interventions to extend the length of pregnancy.

Currently, few studies have focused on the second-trimester plasma lipid profiling of PTB subtypes. Hence, we carried out untargeted lipid profiling of plasma from 258 second-trimester pregnant women, including 44 with extremely or very PTB, 54 with moderate PTB, 71 with late PTB, and 89 with full-term delivery. Reproducible, robust and reliable data were generated and available on MetaboLights, providing a valuable validation dataset for new findings and a test dataset for novel algorithms.

## Methods

### Preterm birth sample recruit

Pregnant women who underwent non-invasive prenatal tests (NIPT) and were willing to donate their samples for research were candidates for this study. Information of pregnancy, delivery and the baby was collected through telephone interview surveys, and samples meeting the following conditions were recruited: 1) singleton; 2) sampling between 13-26 gestational weeks; 3) live birth; 4) vaginal delivery; 5) no pregnancy complications; 6) no newborn anomalies. Preterm birth subtypes were identified as follows: 24w-<32w as extremely or very preterm birth, 32w-<34w as moderate preterm birth, and 34w-<37w as late preterm birth. Women giving birth after 37 complete gestational weeks and with comparable sampling gestational age, maternal age, and baby gender were included as full-term birth. As a result, 44 extremely or very PTB, 54 moderate PTB, 71 late PTB, and 89 full-term delivery women were recruited. For this study, ethical approval was provided by the Institutional Review Board of BGI (FT 15142).

### Plasma sample collection and sample management

Peripheral blood of pregnant women was taken into 5mL EDTA tubes, and plasma was separated through 2-step centrifugation. The blood tube was centrifuged for 10 min at 1,600g at 4 °C, and plasma was transferred and centrifuged for 10 min at 16,000g at 4 °C. The supernatant was isolated and stored at −80°C. A 600 μ L plasma sample each woman was used in the following procedure.

### Extraction of lipids from the blood samples

The lipid extraction was performed following a previously published protocol ^15^. Briefly, plasma samples stored at −80 °C were transferred to −20°C and kept for 30 minutes, thawed at 4 °C and placed on ice. Samples were randomized in order and vortexed for 5s to be homogenized. A 100 μL of each sample was aliquoted into an Eppendorf tube followed by 300 μL of ice-cold isopropanol. Samples were vortexed to mix for 1 minute and placed for 10 minutes at room temperature followed by rested overnight at −20 °C to deposit, after which they were centrifuged to remove the proteins (14000 g, 20 min, 4 °C). A 20 μL supernatant from each sample was transferred into new Eppendorf tubes and 180 μL 50% Methanol was added into each tube and vortexed for 1 minute. The samples were re-randomized and a 20 μL of all the test samples were pooled as a quality control sample (QC sample), which were used for the LC-MS system conditioning before acquisition of the test samples and the quality assessment of the generated data ^16^. The QC and test samples were stored at −80 °C until further analysis.

### UPLC-QTOF MS analysis

The UPLC system used was an ACQUITY UPLC (Waters, USA) and separations were performed on an Acquity UPLC CSH C18 column (2.1×100 mm, 1.7 μm, Waters), with a flow rate of 0.4 mL/min, and a column temperature of 35 °C throughout the experiment. A aliquot of 10 μL samples was injected for analysis, and the lipids were spearated with a gradient mobile phase A comprising of 10 mM ammonium formate with 0.1% formic acid (CAN:H2O = 60:40, v/v) and mobile phase B consisting of 10 mM ammonium formate with 0.1% formic acid (IPA:CAN = 90:10, v/v). The following gradient elution program was adopted for chromatographic separation: 0~2 min, 40-43% phase B; 2.1~6 min, 50-54% phase B; 6.1-8 min, 70-99% phase B; 8.1-10 min, 40% phase B.

Data were acquired on a XEVO-G2XS QTOF mass spectrometer (Waters, UK) equipped with an ion source of electrospray ionization (ESI). The source temperature was 120°C, the desolvation temperature and gas flow were kept at 500°C and 800 L/h. The capillary voltages were set at 2 kV and the sampling cone voltages was 30 V. Scanning was performed in Centroid MSE mode in the mass range 50-2,000 m/z in both positive and negative iron mode with a scan time of 0.2 s. For the MS/MS detection, all fragment ions were generated with a collision energy of 19.0-45.0 eV with a scan time of 0.2 s. The online mass calibration was performed with Leucine enkephalin (LE), which was acquired every 3 seconds during the acquisition. Sample injection order was randomized and distributed into two batches. For each batch, QC samples were analysed ten times at the beginning and three times at the end to equilibrate the column. During acquisition, QC samples were also analysed every ten samples.

### LC-MS data processing

The peak picking and annotation of the MS features were performed on the progenesis QI 2.1 (Waters, Nonlinear Dynamics, USA. The peak picking procedure includes raw file import, peak detection, peak alignment, peak list deisotoping and grouping of adducts (Fig. 1). For the peak detection, the minimal peak intensity was set to be 5000, and the noise ratio and peak width were set to auto in progenesis QI, which automatic optimized during peak detection. After evaluation of the stability of all the samples, the 20150330_POS_QC83 was chosen as a reference sample in positive mode and 20150406_NEG_QC33 in negative mode. All the rest samples were aligned to these two samples in retention time. For the peak grouping, the following adducts were considered, positive mode: [M + Na]^+^, [M + K]^+^, [M + H]^+^ and [M + H – H_2_O]^+^; negative mode: [M - H]^−^ and [M - Cl]^−^.

**Figure 1.**
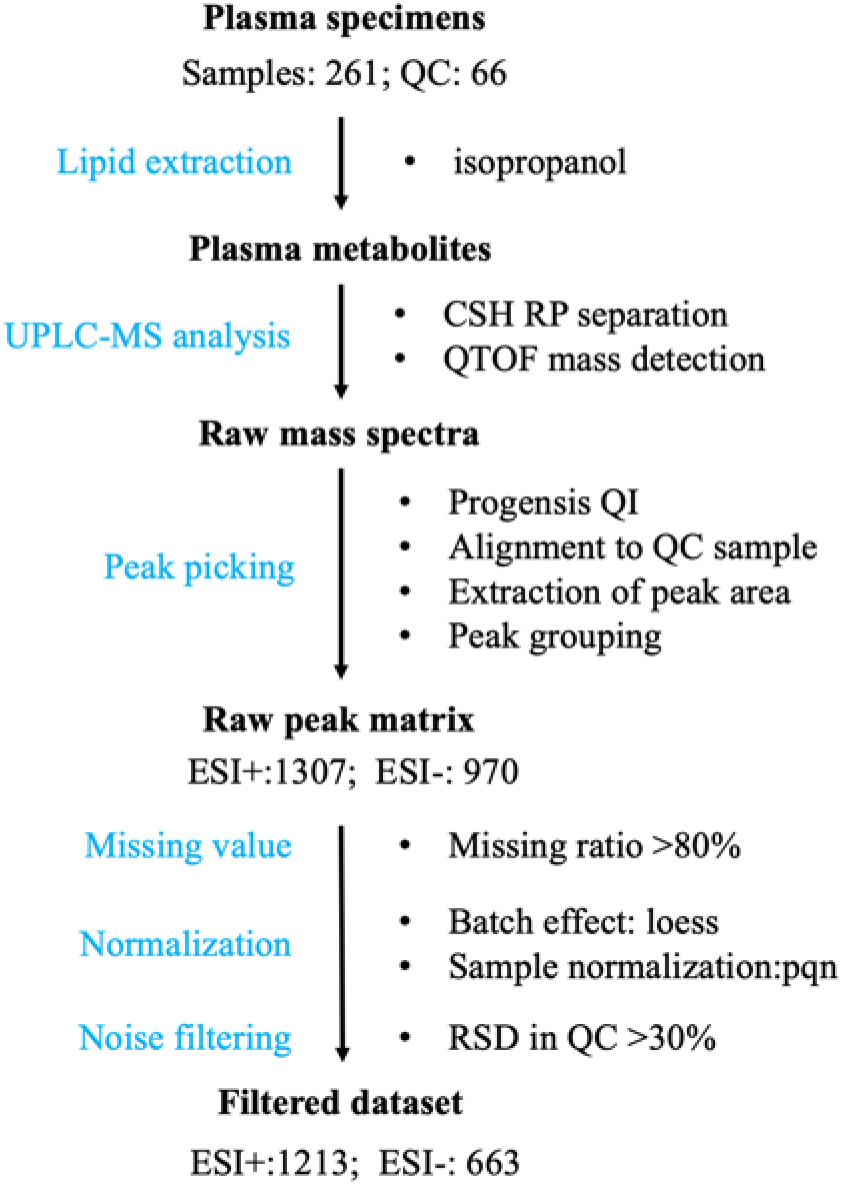
Workflow of the data generation and processing

A data matrix contains m/z paired retention times and peaks intensity was acquired after progenesis QI processing, and the data matrix was further used for data quality assessment using metaX ^17^ (Fig.1). First, features were filtered when their nonzero values are less than 50% of QC samples or removed if they detected in <80% of experimental samples to avoid exogenous impurity. After the filtering, the retained missing values were imputed by KNN ^18^. Then the QC-RLSC ^19^ method was applied to correct each metabolic feature signal draft according to the signal change in QC samples and the PQN ^20^ was performed to normalize all the samples. The features with RSD >30% in QC samples were removed after the QC-RLSC and PQN correction to get clean data, in which unwanted variations were removed.

### Putative annotation

The putative annotation of features was conducted by searching against online databases using accurate mass, isotope abundance and MS/MS fragmentation pattern using Progenesis QI. The initial tentative identifications were obtained by searching the accurate mass of the MS features against the following databases within a mass error (10ppm): Human Metabolome database (http://www.hmdb.ca/), Kyoto Encyclopedia of Genes and Genomes (https://www.genome.jp/kegg/) and LipidMaps (http://www.lipidmaps.org/). The theoretical isotope peak distribution of the candidates was calculated and compared with that of the MS features to further filter the tentative identifications. MS/MS fragmentation matching pattern was also evaluated between the theoretical MS/MS spectra and the deconvoluted experimental MS/MS spectra with a mass error of10 ppm. A putative identification list with annotation scores was acquired after the annotation process.

### Statistical analysis

Principal component analysis and relative standards deviations, which were employed for the quality assessment were performed using the R version 3.4.0.

### Data Records

The raw data files in the .raw format generated from untargeted UPLC-MS profiling in both the positive and negative mode are submitted to the MetaboLights repository (Data Citation 1:MetaboLights MTBLS885). The peak picking and annotation obtained from Progenesis QI 2.1 data processing were also contained in the deposited data, and a detailed explanation to the headers was provided within these tables. Furthermore, the raw data files and the processed results have also been submitted to CNSA (Data Citation 2: CNP0000369), researchers may download the raw data via FTP: https://db.cngb.org/. Raw data files and the corresponding raw peak tables, the processed peak tables and scripts are listed in Table 1.

**Table 1.** Description of the data records

| Type | Description | File Name | Location |
| --- | --- | --- | --- |
| Raw data | Raw data file in .raw format | *.raw | Metabolights (MTBLS885) & CNSA (CNP0000369) |
| Raw data matrix | Raw data matrix generated from peak picking using Progenesis QI 2.1 | raw_peak_table_neg.csv | Metabolights (MTBLS885) & CNSA (CNP0000369) |
|  |  | raw_peak_table_pos.csv |  |
| Sample list | Sample information used as input for processing and quality validation script | sample_list_neg.txt |  |
|  |  | sample_list_pos.txt |  |
| Scripts | Scripts used for data processing (missing value processing, batch effect correction, sample normalization and noise filtering) and quality validation (PCA and CV plots) | data_process_and_quality_validation_neg.R |  |
|  |  | data_process_and_quality_validation_pos.R |  |
| Processed data matrix | Clean data matrix after data processing | processed_peak_table_neg.csv |  |
|  |  | processed_peak_table_pos.csv |  |
| Putative identification | Putative identification list | putative_identification_neg.xlsx |  |
|  |  | putative_identification_pos.xlsx |  |

### Technical Validation

The quality validation of lipids profiling was performed following the best practices of analytical methodologies ^21,22^. All the plasma samples were analysed in a random order, and the stability and reproducibility were evaluated by the pooled QC samples inserted into the analysis sequence during the whole experimental period, as shown in Fig. 2. The data quality was measured in the following three ways: (1) the overall signal drift indicated by overlap of the total ion chromatograms (TIC) plot of quality control samples, (2) the reproducibility measured by the relative standard deviation of all the features from the quality control samples and (3) the overall variance represented by PCA plots. The TIC plots of all the pooled quality control samples in positive and negative mode (Fig. 2 A, B) showed that these quality control samples were highly overlapped without obvious retention time and intensity drift, indicating a good signal consistence throughout the data acquisition. A total of 1306 MS features were detected in the positive mode and 734 in the negative mode. After removing severe missing features, 1287 and 730 features in positive and negative mode were remained. After batch effect correction, sample normalization, the ratio of MS features in QC samples with relative standard deviation ≤ 30% accounts for 92.57% and 91.01% respectively in the positive and negative mode (Fig2. C, D), indicating the data acquisition and processing resulted into a highly reproducible dataset. The principal component analysis (PCA) was performed among the test samples and the QC samples to assess the experimental variance. The pooled QC samples were tightly clustered within the biological samples in both positive and negative (Fig. 2E, F), which was consistent with our expectation that pooled samples are most similar with each other, indicating the analysis process met the required qualifications.

**Figure 2.**
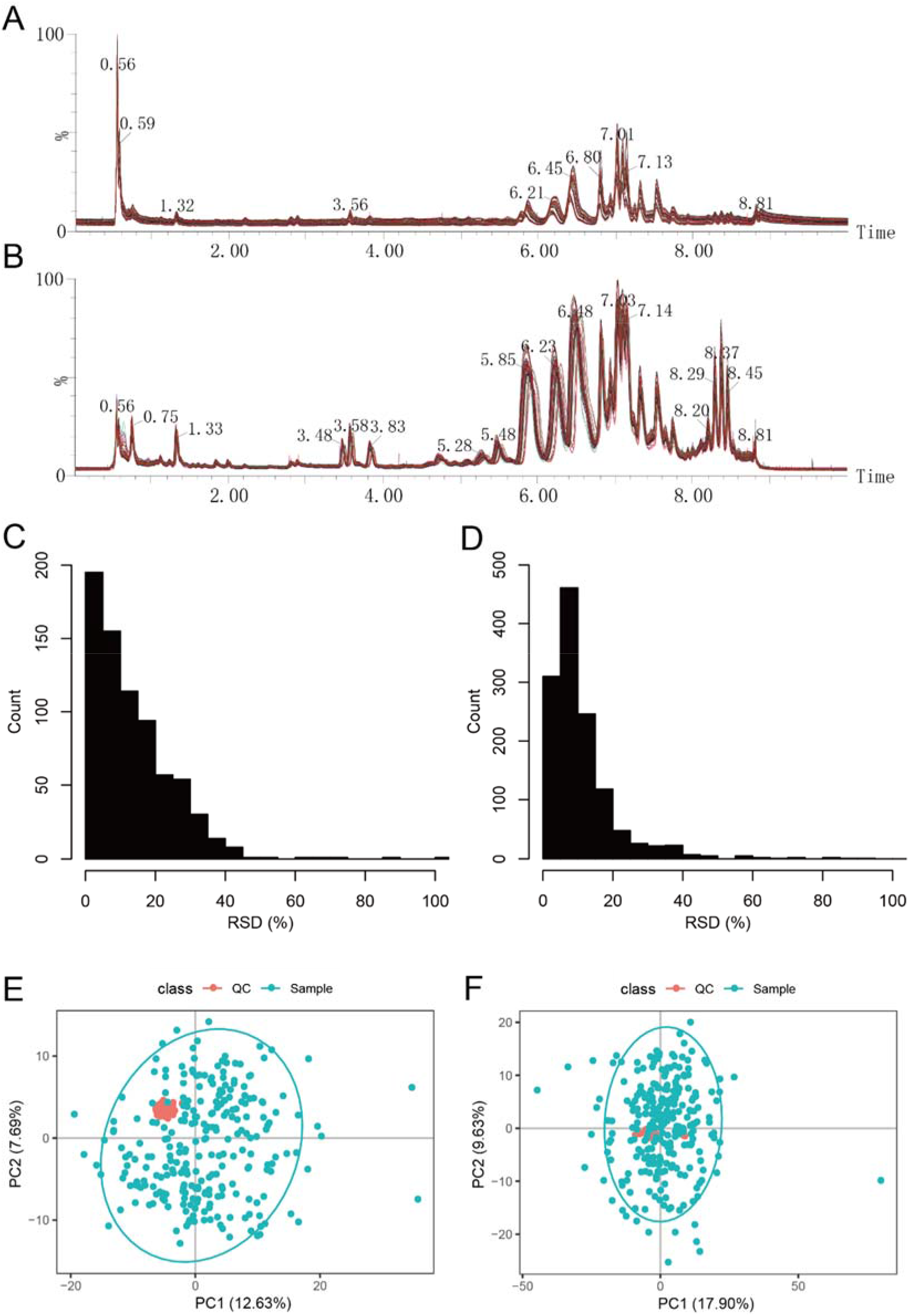
Data quality assessment. Total Ion Chromatography of the negative (A) and positive mode (B); Reproductivity of the quality control samples in the negative (C) and positive mode (D); PCA plots of the negative (E) and positive mode (F).

### Usage Notes

The raw data files in .raw format, the raw peak matrix, clean peak matrix, the putative identification list and a detailed explanation to the header of these files are available from MetaboLights (Data Citation 1) and the CNSA (Data Citation 2). Since the raw data were generated from MSE mode, a branch of the data independent acquisition, the MS-DIAL ^23^ and Progenesis QI, were recommended to redo the peak picking and identification. The instrument .raw files could be accessed directly using MassLynx or converted into .mzXML format using databridge and checked by the Mzmine ^24^. The raw peak matrix files were generated from the raw files using Progensis QI, and gone through removing severe missing peaks, missing value imputation, batch effect correction, sample normalization and noise filtering to generate the clean peak matrix, which was further used for data quality validation using metaX. All the codes used in data cleaning were also uploaded to Metabolights. The clean peak matrix is assumed to be free of unwanted variation but still need to be log-transformed to correct the non-normal distribution before univariate or multivariate analysis. All together with the identification list, this dataset may be used as a reference dataset for the evaluation of potential biomarkers discovered by other studies, as well as validation evidence of the findings of other omics datasets. The raw spectra and the raw data matrix may contribute to the evaluation of novel spectra processing software and new algorithms/pipeline for data preprocessing.

## Data Citations

1. Kang Huang et al. MetaboLights MTBLS885 (2019)
2. CNSA https://db.cngb.org/cnsa: accession number CNP0000369.

## Acknowledgements

We are grateful to the participants who donated their samples for the study. This project is supported by the National Natural Science Foundation of China (No.81300075), the Natural Science Foundation of Guangdong Province (No. 2014A030313795), the Shenzhen Municipal Government of China (No.JCYJ20170412152854656, JCYJ20180703093402288).

## Author Information

W.J.W. and H.J. conceived the study. L.Y., M.C., Y.C., S.Z. and J.Z. performed the assays. X.Y. and X.C. performed the LC-MS data acquisition. Z.M. and K.H. performed the LC-MS data processing. Z.M. and K.H. prepared the figures and tables. Z.M., W.J.W., L.Y., & H.J. wrote the manuscript.

